# Anionic phospholipids determine G protein coupling selectivity at distinct subcellular membrane compartments

**DOI:** 10.1101/2023.08.31.555797

**Authors:** Evelyn H. Hernandez, Aashish Manglik, Roshanak Irannejad

## Abstract

G protein-coupled receptors (GPCRs) activate different signaling pathways through coupling to heterotrimeric G proteins consisting of four types of G alpha (Gα) subunits and the beta gamma (ýy) subunits. The C-terminal regions of Gα proteins is thought to determine coupling selectivity. However, there are several reports indicating that some receptors are promiscuous and can couple to multiple Gα proteins. The precise mechanism promoting promiscuous coupling is not fully understood. Here we show that anionic phospholipids such as phosphoinositide 4,5 bisphosphate (PI(4,5)P2) promote electrostatic interactions between anionic lipid head groups and positively charged amino acids in the transmembrane 4 (TM4) region of beta adrenergic receptors (βARs) to prime receptor coupling to the cognate Gαs and non-cognate Gαi proteins. The promiscuous coupling can only occur at the plasma membrane, a membrane compartment enriched in PI(4,5)P2, while receptors only couple to cognate Gαs proteins in compartments that are enriched in less negatively charged phosphoinositides such PI4P at the Golgi membranes. We took advantage of the rapamycin dimerization system to rapidly and inducibly deplete PI(4,5)P2 by recruiting a phosphatase to the plasma membrane and found that in the absence of anionic phospholipids, βARs couple only to their cognate Gαs protein. Finally, we found that mutating βARs PI(4,5)P2 binding motifs or depleting PI(4,5)P2 results in enhanced βARs-mediated cAMP response. Together, these findings reveal a role for anionic phosphoinositides in regulating GPCR activity at different subcellular compartments.

## INTRODUCTION

There are over 800 types of GPCRs that relay their signals by coupling to sixteen types of Gα proteins, classified into four different families of Gαs, Gαi/o, Gαq/11 and Gα12/13^1^. How coupling selectivity is precisely determined is still unclear. The C-terminal domains of Gα proteins have been classically attributed to determine Gα protein coupling selectivity. The C-terminal domains of different Gα protein types are distinct especially in the last ten amino acids. This is the region that embeds itself into the core of the receptor in the open active conformation ^2–5^. While the significance of the Gα protein C-terminal domain for coupling selectivity has been extensively studied, more recent evidence highlights the importance of the N-terminal region of the Gα protein ^4, 6–8^. By analyzing the coupling efficiency of various types of Gα protein chimeras that contained a combination of C-and N-terminal domains of Gαs, Gαi, Gαq and Gα12, it was shown that while the C-terminal domain plays a significant role in determining the coupling selectivity, the N-terminal core region of the Gα proteins also contributed to determining selectivity (Figure 1a) ^7^.

**Figure 1.**
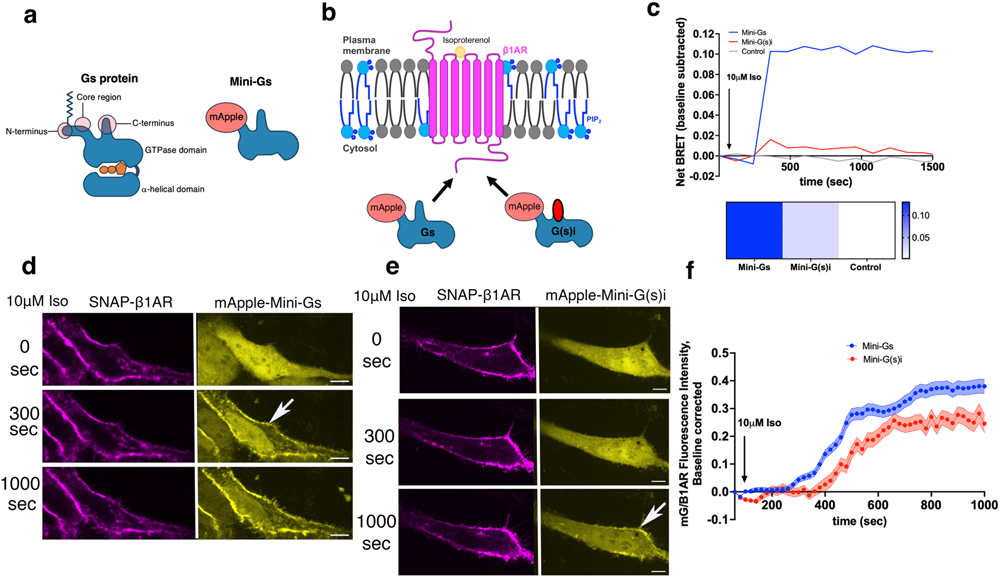
β1AR couples to Gαs and Gαi proteins at the plasma membrane upon isoproterenol (Iso) stimulation, as detected by Mini-Gs and Mini-G(s)i recruitment. **(a)** Schematic representation of G protein highlighting the specific domains compared to Mini-Gs. (**b**) Schematic representation of Mini-Gs and Mini-G(s)i coupling to β1AR at the plasma membrane. Red oval represents Gi C-terminal sequence. (**c**) Time-course (top) and heat-map (bottom) representation of BRET analysis of Flag-β1AR-NanoLuc with mVenus tagged Mini-Gs and Mini-G(s)i. Each value represents the mean normalized ΔBRET over 20 minutes. N= 3 biological replicates. (**d**) Confocal images of representative SNAP-tagged β1AR-expressing HeLa cells with mApple-Mini-Gs and (**e**) mApple-Mini-G(s)i expression before and after 10 μM Iso addition. Stimulation with 10 μM Iso results in the recruitment of Mini-Gs (n= 38 cells, 3 biological replicates) and Mini-G(s)i (n= 26 cells, 5 biological replicates) to active β1AR at the plasma membrane in HeLa cells. Arrows indicate recruited Mini-Gs and Mini-G(s)i at plasma membrane; Scale bar = 10 μm. (**f**) Quantification of Mini-Gα recruitments to activated β1AR at the plasma membrane in HeLa cells after 10 μM Iso addition; normalized fluorescence intensity of Mini-G probes at plasma membrane relative to SNAP-tagged-β1AR. Quantifications were baseline corrected.

A wide range of, *in vitro,* biochemical, and structural analyses began to reveal the molecular mechanisms determining GPCR-Gα protein coupling selectivity. A comprehensive analysis of GPCR and Gα protein sequences unveiled a ‘selectivity barcode’ that dictates Gα protein coupling patterns ^4^. While a specific amino acid sequence for a selectivity barcode was not determined, amino acid properties such as electrostatic charge, hydrophobicity, and side chain size were found to be the determining factor in GPCR-Gα protein binding interfaces. In addition, post-translational modifications, phosphorylation, and potential allostery provided by membrane phospholipids or phosphoinositides were indicated to determine selectivity ^4^. A more recent study found that the intracellular loop 3 regions (ICL3) of GPCRs are allosterically modulated to allow access to the GPCR core for Gα proteins coupling ^9^.

As integral membrane proteins, GPCRs are intimately associated with lipids, but the role of lipids in protein function and signaling is poorly understood. *In vitro* studies suggest that membrane lipid compositions may play a role in determining Gα protein coupling selectivity ^10–12^. In particular, anionic phospholipids such as phosphoinositide 4,5 bisphosphate (PI(4,5)P2), which are highly enriched at the plasma membrane, promote Gαs coupling via electrostatic interactions between anionic lipid head groups and positively charged amino acids in the core region of Gαs ^8^. Other groups have demonstrated that anionic phospholipids provide a priming mechanism for the receptor to engage signaling partners ^12^. Phospholipids were shown to embed themselves into the G protein binding site and stabilize the receptor in an open active conformation to facilitate Gα protein coupling ^12^. This mechanism suggests that negatively charged phospholipids directly prepare GPCRs to engage intracellular Gα proteins. This is reminiscent to previous models suggesting that non-cognate Gα proteins could synergistically prime GPCRs to bind to cognate Gα protein subtypes and enhance the signaling effect ^13^.

While many of these studies have focused on coupling selectivity, there are also several studies that suggest some receptors can couple to non-cognate Gα proteins in addition to their cognate Gα proteins. For example, β2 adrenergic receptor (β2AR) and muscarinic receptor 4 have been reported to couple to multiple Gα proteins with various efficiencies ^14, 15^ serotonin type 4 receptor couples to Gαs and Gαi proteins ^16^, and the Neurokinin-1 receptor couples to Gαs and Gαq^17^. Although β1AR coupling to Gαs protein has been widely studied, there is evidence that it can also be Gαi-coupled, where inhibition of Gαi signaling by pertussis toxin influenced β1AR-mediated signaling^14, 18^. What promotes the promiscuous coupling is not fully understood. A recent study suggests that most GPCRs interact with all four families of nucleotide-decoupled G proteins, however, these interactions do not necessarily lead to GDP release of the noncognate Gα proteins and they retain a preference for cognate G proteins^15^.

In the current study, we investigate the role of phospholipids in regulating coupling selectivity of an exemplary receptor, β1AR, to two opposing Gα proteins. We found that PI(4,5)P2 at the plasma membrane plays a role in priming the receptors, allowing for both Gαs and Gαi protein coupling, while β1AR can only couple to Gαs at the Golgi membranes that are enriched in PI(4)P. We found that mutating the positively charged residues of β1AR at TM4 that have been reported to interact with PI(4,5)P2 ^11^ disrupts this dual coupling and biases receptor coupling to Gαs only. Similarly, depleting anionic phospholipids such as PI(4,5)P2, by rapidly recruiting a specific phosphatase at the plasma membrane using the rapamycin dimerization strategy, results in β1AR coupling only to its cognate Gαs protein. This dual coupling mechanism is not unique to β1AR, as we have also observed it for another receptor with similar PI(4,5)P2 binding motifs such as β2AR. Finally, we demonstrate that isoproterenol-mediated cAMP production is enhanced in cells expressing β1AR PI(4,5)P2 binding mutants or when PI(4,5)P2 is depleted. Altogether, these results suggest that PI(4,5)P2 plays a role in priming the active conformation of receptors and facilitating promiscuous Gα protein coupling to β1AR and β2AR at the plasma membrane. These findings also provide evidence for the role of phosphoinositides in regulating GPCR coupling selectivity at different subcellular compartments.

## RESULTS

### β1AR couples to Gαs and Gαi proteins with distinct efficiency and kinetics

Several studies suggest that GPCRs can couple to multiple Gα subunits. What contributes to this promiscuous coupling is not fully understood. Recent evidence suggest that the C-terminal domains of Gα proteins are not the only determinant sequence for coupling selectivity and the core region of Gα proteins may also play a role in promoting GPCR-G protein coupling ^4, 7, 19^. This has been shown particularly for Gαs involving the electrostatic interactions between the positively charged residues on the intracellular loops of GPCRs with anionic phospholipids such as PI(4,5)P2 and the positively charged amino acids in the core region of Gαs ^11, 19^. To begin to test the role of these electrostatic interactions, we first focused on the β1AR because it not only couples to its cognate Gαs protein, but evidence suggests that β1AR-mediated cAMP response is sensitive to pertussis toxin, indicating involvement of Gαi coupling^14, 18^. To assess whether β1AR can couple to both Gαs and Gαi, we employed Mini-G proteins, which are engineered GTPase Ras-like domains of Gα proteins that conserve the associations formed between activated GPCR and its respective Gα protein^20 21^. These Mini-G proteins are designed to only interact with activate conformations of the GPCR and are found to uncouple nucleotide binding and receptor association. The N-terminal region of the G proteins which contains lipid modifications required for membrane anchoring are also absent in the Mini-G proteins (Figure 1a)^21, 22^. Due to stability issues during the purification process for structural studies, the Mini-G tool used to represent Gαi protein was made as a chimera of Gαs and Gαi sequences containing the C-terminus of Gαi and the core regions of Gαs (Figure 1b), thus we refer to this tool as Mini-G(s)i ^21^. To monitor the interaction of Mini-G proteins to activated β1AR, we first used bioluminescence resonance energy transfer (BRET) in HeLa cells expressing β1AR fused to NanoLuc luciferase (Flag-β1AR-NanoLuc) with mVenus tagged to either Mini-Gs and Mini-G(s)i at the N-terminal.

Stimulation of HeLa cells with 10 μM isoproterenol resulted in a pronounced increased in energy transfer between β1AR and Mini-Gs. Interestingly, we also observed BRET between β1AR and Mini-G(s)i, although less efficiently (Figure 1c). These data suggest that β1AR engages the C-terminal domains of both Gαs and Gαi as detected by Mini-Gs and Mini-G(S)i probes.

To better monitor the cellular location where this dual coupling is occurring, we then expressed SNAP-tagged β1AR and Mini-Gs fused to mApple (mApple-Mini-Gs) in HeLa cells. In unstimulated cells, mApple-Mini-Gs was diffuse throughout the cytoplasm (Figure 1d, top panels; 0 min). Upon stimulation of cells with 10 μM Isoproterenol, a βAR full agonist, mApple-Mini-Gs was recruited to the plasma membrane starting at ∼300 seconds (Figure 1d, top panel and 1f, blue line). HeLa cells expressing SNAP-tagged β1AR and Mini-G(s)i fused to mApple (mApple-Mini-G(s)i), also showed recruitment to the plasma membrane, however, at a later time point, starting at ∼400 seconds (Figure 1e, bottom panels and 1f, red line).

Importantly, this dual coupling was only observed at the plasma membrane when cells were stimulated with membrane impermeant full agonist, isoproterenol. We have previously shown that β1AR activates Gαs protein-mediated signaling from the plasma membrane as well as the Golgi ^23–25^. To monitor whether this dual coupling can occur at the Golgi, HeLa cells were stimulated with epinephrine, another full agonist of βARs that is able to reach Golgi-localized β1AR. Epinephrine is a substrate for an organic cation transporter (OCT3), therefore it can be transported to the Golgi membranes^23, 24, 26^. Stimulating β1AR expressing HeLa cells with 10 μM epinephrine resulted in the recruitment of both mApple-Mini-Gs and mApple-Mini-G(s)i to the plasma membrane like what we observed with isoproterenol. However, epinephrine stimulation resulted in the recruitment of mApple-Mini-Gs only to Golgi-localized β1AR (Supplementary Figure 1a and b). When HeLa cells were stimulated with dobutamine, a β1AR partial agonist that can also reach the Golgi membrane due to its hydrophobicity, we found that coupling was biased towards Mini-Gs only, at both the plasma as well as the Golgi membranes (Supplementary Figure 2). Altogether, these data suggest that full agonists (epinephrine and isoproterenol) can prime β1AR at the plasma membrane, allowing for coupling of both the cognate and non-cognate Gα proteins, while dobutamine, a partial agonist, biases this coupling toward the cognate G protein, Gαs. Moreover, this dual coupling is only observed at the plasma membrane, a membrane compartment enriched in negatively charged phospholipids such as PI(4,5)P2 and not at the Golgi membranes, which are enriched in PI(4)P ^27^, suggesting that the lipid composition of subcellular membrane compartments may play a role in modulating β1AR-Gα protein coupling promiscuity.

### Positively charged residues at the intracellular TM4 region of β1AR promote dual Gαs and Gαi coupling

The lack of promiscuous coupling of Mini-Gs and Mini-G(s)i to activated β1AR at the Golgi membranes (Supplementary Figure 1) suggested the involvement of negatively charged phosphoinositides such as PI(4,5)P2 in determining β1AR-Gα dual coupling. Anionic phospholipids have been shown to facilitate GPCR-G protein coupling and signaling through bridging interactions between cationic residues on receptor and Gαs protein surfaces and directing allostery of GPCR activity ^10, 11, 19, 28^. Certain class A GPCRs that preferentially couple to Gαs interact with anionic phospholipids PI(4,5)P2, which are highly enriched at plasma membranes, over other phosphoinositides such as PI(3)P and PI(4)P that are found at the endosomes and Golgi membranes, respectively ^11, 27^. The C-terminal end of N-terminal region of Gαs, where we refer to as the “core region” (Figure 1a) contains positively charges residues that have been shown to form electrostatic interactions with anionic phospholipids and the positively charged residues on the intracellular loops of class A GPCRs ^19^ (Supplementary Figure 3a). The mApple-Mini-G(s)i is a chimera of the core region of Gαs and the C-terminus of Gαi protein (Supplementary Figure 3a and 4). Thus, to assess the effect of these positively charged residues (KDKQ) on the recruitment of mApple-Mini-G(s)i chimera to activated β1AR at the plasma membrane, we mutated the positively charged residues (KDKQ) to the negative residues (EDGE) that are present in Gαi proteins ^29^ (Supplementary Figure 3a and 4). Stimulating HeLa cells with isoproterenol resulted in mApple-Mini-G(s)i EDGE recruitment to β1AR at the plasma membrane, however with less efficiency and starting ∼500 seconds (Supplementary Figure 3b and d). Interestingly, this weak recruitment at the plasma membrane was significantly enhanced when mG(s)i EDGE C-terminus was swapped with Gαs C-terminus (Mini-G(s)i EDGE w/ Gs CTD) (supplementary Figure 3c and d and 4), suggesting that the core and the C-terminal regions of Gα proteins are both involved in coupling to β1AR. Importantly, the positively charged residues on the core region of Gαs protein contribute to faster and more efficient coupling of the cognate Gαs protein to activated β1AR at the plasma membrane.

PI(4,5)P2 binding sites were identified for a variety of Gαs coupled class A GPCRs such as neurotensin receptor 1 (NTSR1) and turkey β1AR ^11^. These binding sites include primarily amino acid residues with positive charges such as lysine (K) and arginine (R). The sites that provided the largest disruption to phosphoinositide interaction were enriched in the TM4 region of NTSR1 ^11^. To identify similar amino acids in human β1AR, we aligned human β1AR sequence with NTSR1 and turkey β1AR and found the corresponding and homologous residues (Supplementary Figure 5). We mutated these positively charged amino acids in the TM4 of SNAP-tagged human β1AR (RARAR) to neutral residues (IATAL) (Figure 2a). To assess the consequence of disrupting the electrostatic interactions between the receptor and PI(4,5)P2 in coupling promiscuity, we compared Mini-Gs and Mini-G(s)i recruitments to β1AR IATAL mutant. We matched the expression level of wild type β1AR and β1AR IATAL mutant (Supplementary Figure 6a). We found that β1AR IATAL mutant impedes recruitment of Mini-G(s)i to the plasma membrane (Figure 2b-d). These data suggest that the electrostatic interaction between the positively charged residues at TM4 region of β1AR and PI(4,5)P2, primes the receptor for dual coupling to Mini-Gs and Mini-G(s)i at the plasma membrane.

**Figure 2.**
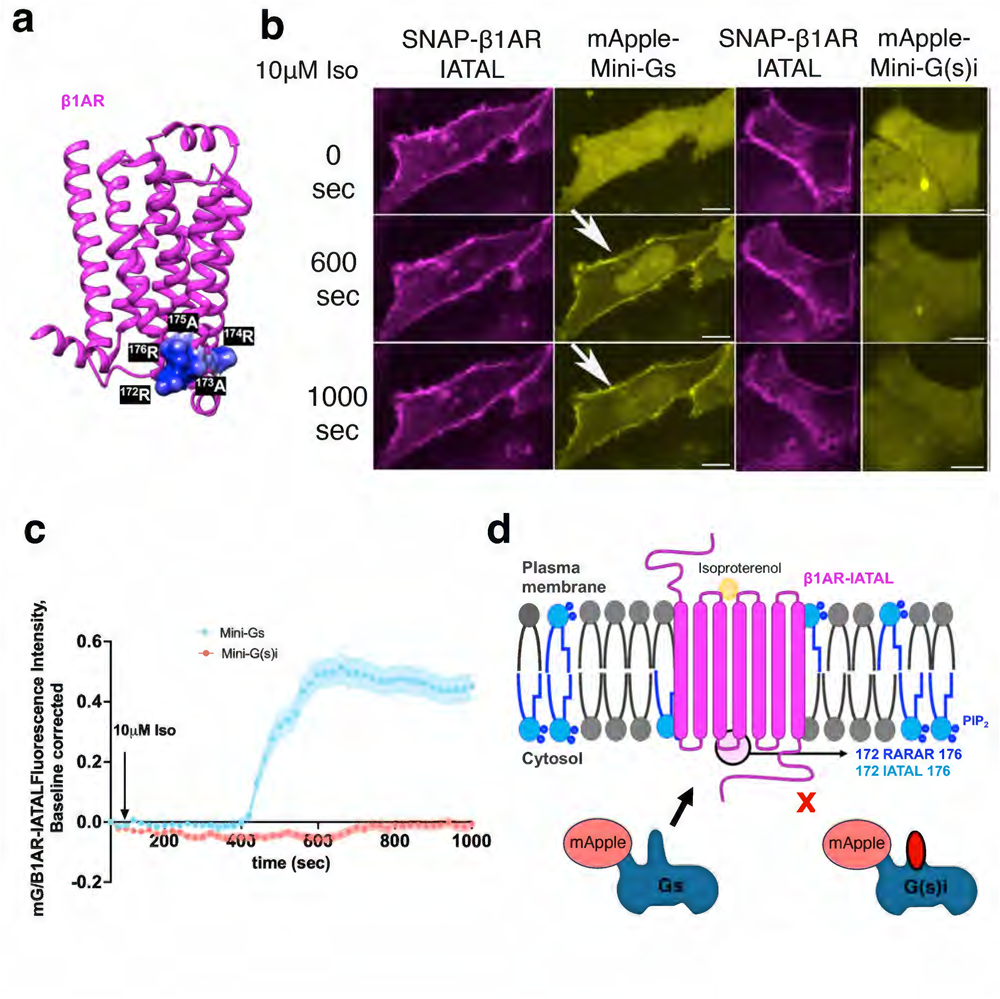
β1AR PI(4,5)P2 binding mutant cannot recruit Mini-G(s)i. (**a**) β1AR crystal structure (PDB: 7BTS) with predicted PI(4,5)P2 binding motif at residues 172-176 (RARAR) showing positive electrostatic charge. (**b**) Confocal images of representative SNAP-tagged mutant β1AR IATAL-expressing HeLa cells (magenta panels) with mApple-Mini-Gs and mApple-Mini-G(s)i (yellow panels) expression before and after 10 μM Iso addition. Stimulation with 10 μM Iso results in recruitment of Mini-Gs but not Mini-G(s)i to active mutant β1AR IATAL at the plasma membrane in HeLa cells (Mini-Gs: n= 44 cells, 4 biological replicates; Mini-G(s)i: n= 30 cells, 4 biological replicates). Arrow indicates recruited Mini-Gs at plasma membrane; Scale bar = 10 μm. (**c**) Quantification of Mini-G recruitments to activated β1AR IATAL at the plasma membrane in HeLa cells after 10 μM Iso addition; normalized fluorescence intensity of Mini-G probes at plasma membrane relative to SNAP-tagged-β1AR. Quantifications were baseline corrected. (**d**) Schematic representation of Mini-Gs coupling to mutated β1AR. Residues 172-176 were mutated from their positive RARAR to neutral residues IATAL. Red oval represents Gi C-terminal sequence.

### PI(4,5)P2 contributes to βAR-Gα protein promiscuous coupling at the plasma membrane

To further investigate how PI(4,5)P2 regulates β1AR/Gα protein dual coupling at the plasma membrane, we next depleted PI(4,5)P2 by targeting phosphoinositide phosphatases and testing whether this modification affects Gα protein coupling selectivity. Using the FKBP-FRB rapamycin system, we specifically targeted FKBP to the plasma membrane using a known plasma membrane targeting sequence (first 11 amino acids of Lyn) (PM-FKPB) and cytosolically expressed the FRB fragment fused to mVenus and a chimeric PI(4)P (Sac1) and PI(5)P (INP55E) phosphatase, named Pseudojanin (mVenus-FRB-Pseudojanin) that depletes phosphate 4 and 5 groups from PI(4,5)P2 ^27^ (Figure 3a). Upon addition of 1 μM rapamycin, mVenus-FRB-Pseudojanin was recruited to the plasma membrane, depleting PI(4,5)P2, as detected by a decrease in PH-PLC8 (a PI(4,5)P2 maker) localization on the plasma membrane, (Figure 3a-c). We then expressed PM-FKBP and mVenus-FRB-Pseudojanin to assess the kinetics of Mini-Gs and Mini-G(s)i recruitment to wild-type β1AR (Figure 3d). We pre-incubated these cells with 1 μM rapamycin for 10 min and looked for cells where PI(4,5)P2 were depleted. Upon depletion of PI(4,5)P2, mApple-Mini-Gs was recruited to the activated β1AR at the plasma membrane (Figure 3d, top panel and 3e, green line) while mApple-Mini-G(s)i was not (Figure 3d, bottom panel, 3e orange line). This is consistent with what we observed with β1AR IATAL mutant (Figure 2).

**Figure 3.**
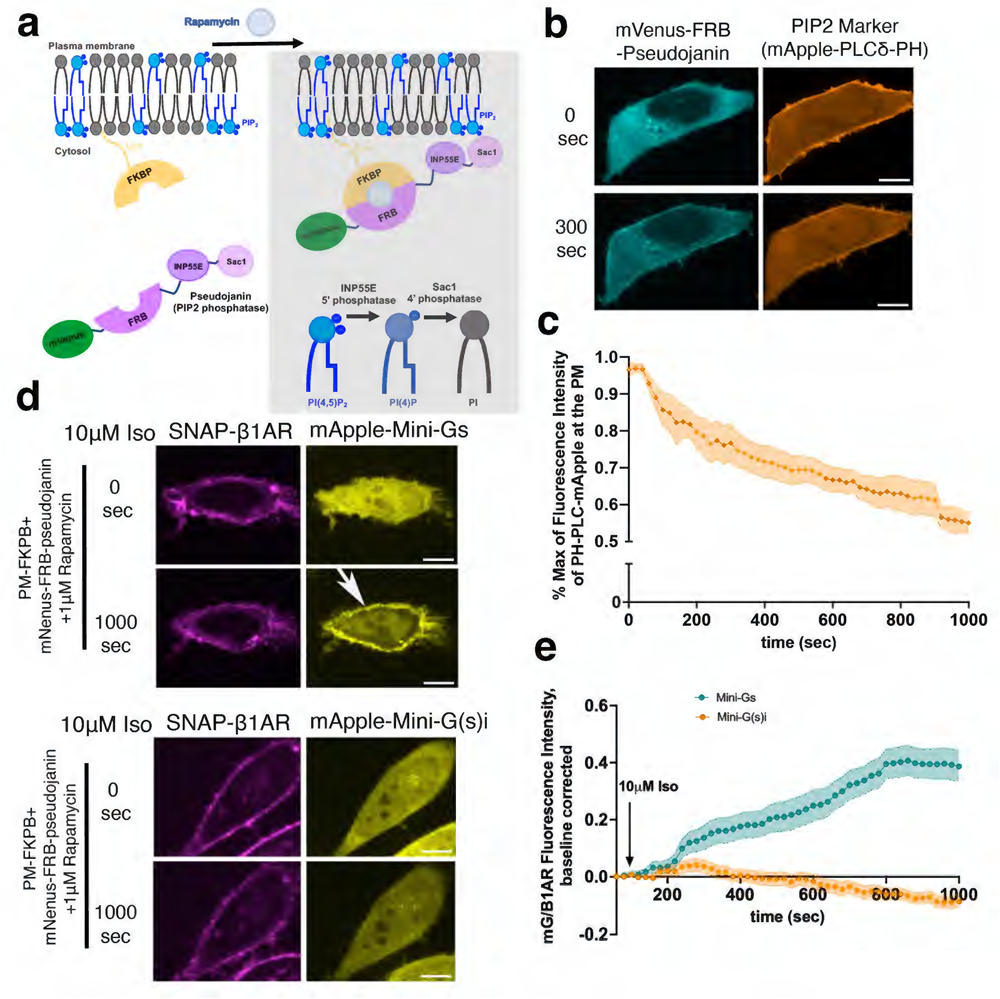
PI(4,5)P2 depletion at the plasma membrane disrupts Mini-G(s)i recruitment to activated β1AR. (**a**) Schematic of FKBP-FRB system where the FKBP fragment is targeted to the plasma membrane using a Lyn motif and the FRB fragment is fused to mVenus and a chimeric phosphatase (Pseudojanin) containing INP55E and Sac1, 5’ and 4’ phosphatases, respectively. Upon the addition of 1 μM Rapamycin, the FKBP and FRB fragments dimerize allowing the chimeric phosphatase access to deplete PI(4,5)P2 at the plasma membrane (**b**) Representative confocal images of PI(4,5)P2 depletion using the FKBP-FRB system described. mVenus-FRB-Pseudojanin is recruited to the plasma membrane upon addition of 1 μM Rapamycin (left panels) and the PI(4,5)P2 marker, an mApple tagged pleckstrin homology (PH) domain (mApple-PLCδ-PH, right panels), is translocated to the cytosol after 300 seconds. (**c**) Quantification of mApple-PLCδ-PH translocation from the plasma membrane upon PI(4,5)P2 depletion after 1 μM Rapamycin treatment; normalized fluorescence intensity of mApple-PLCδ-PH at plasma membrane relative to maximum value. (**d**) Representative confocal images of SNAP-tagged β1AR-expressing HeLa cells after PI(4,5)P2 depletion with mApple-Mini-Gs (top panels) and mApple-Mini-G(s)i (bottom panels) expression before and after 10 μM Iso addition. Stimulation with 10 μM Iso results in the recruitment of Mini-Gs but not Mini-G(s)i to active β1AR at the plasma membrane (Mini-Gs: n= 20 cells, 7 biological replicates; Mini-G(s)i: n= 23 cells, 4 biological replicates). Arrow indicates recruited Mini-Gs at plasma membrane; Scale bar = 10 μm. (**e**) Quantification Mini-Gα recruitments to activated β1AR upon PI(4,5)P2 depletion in HeLa cells after 10 μM Iso addition; normalized fluorescence intensity of Mini-G probes at plasma membrane relative to SNAP-tagged-β1AR. Quantifications were baseline corrected.

To further confirm the role of PI(4,5)P2 in promoting coupling promiscuity, we next focused on β2AR, another closely related adrenergic receptor that has been shown to couple to both Gαs and Gαi ^30, 31, 32^. We first confirmed this dual coupling using mApple-Mini-Gs and mApple-Mini-G(s)i and similarly found faster kinetics in the Mini-Gs coupling followed by Mini-G(s)i coupling in response to 10 μM isoproterenol (Figure 4a-c). To identify similar amino acids that were shown to bind PI(4,5)P2 in NRTS1 and β1AR, we aligned human β1AR sequence with β2AR and found the corresponding and homologous residues (Supplementary Figure 7). We then mutated these residues to neutral residues (INTAL) (Figure 4d). To assess the consequence of disrupting the electrostatic interactions between β2AR and PI(4,5)P2 in coupling promiscuity, we compared Mini-Gs and Mini-G(s)i recruitments to β2AR INTAL mutant. We matched the expression level of wild type β2AR and β2AR INTAL mutant (Supplementary Figure 6b). Similar to what we found for β1AR PI(4,5)P2 binding mutant, β2AR INTAL also impedes recruitment of Mini-G(s)i to the plasma membrane (Supplementary Figure 8a-b, Figure 4e). Depleting PI(4,5)P2 using the Pseudojanin rapamycin system at the plasma membrane resulted in decreased β2AR endocytosis (Supplementary Figure 8c)^33–35^. Nonetheless, depleting PI(4,5)P2 biased β2AR coupling to Gαs at the plasma membrane (Supplementary Figure 8d). Altogether, these data further confirm that PI(4,5)P2 primes the receptor to facilitate dual coupling of Gαs and Gαi at the plasma membrane via electrostatic interactions with the TM4 region.

**Figure 4.**
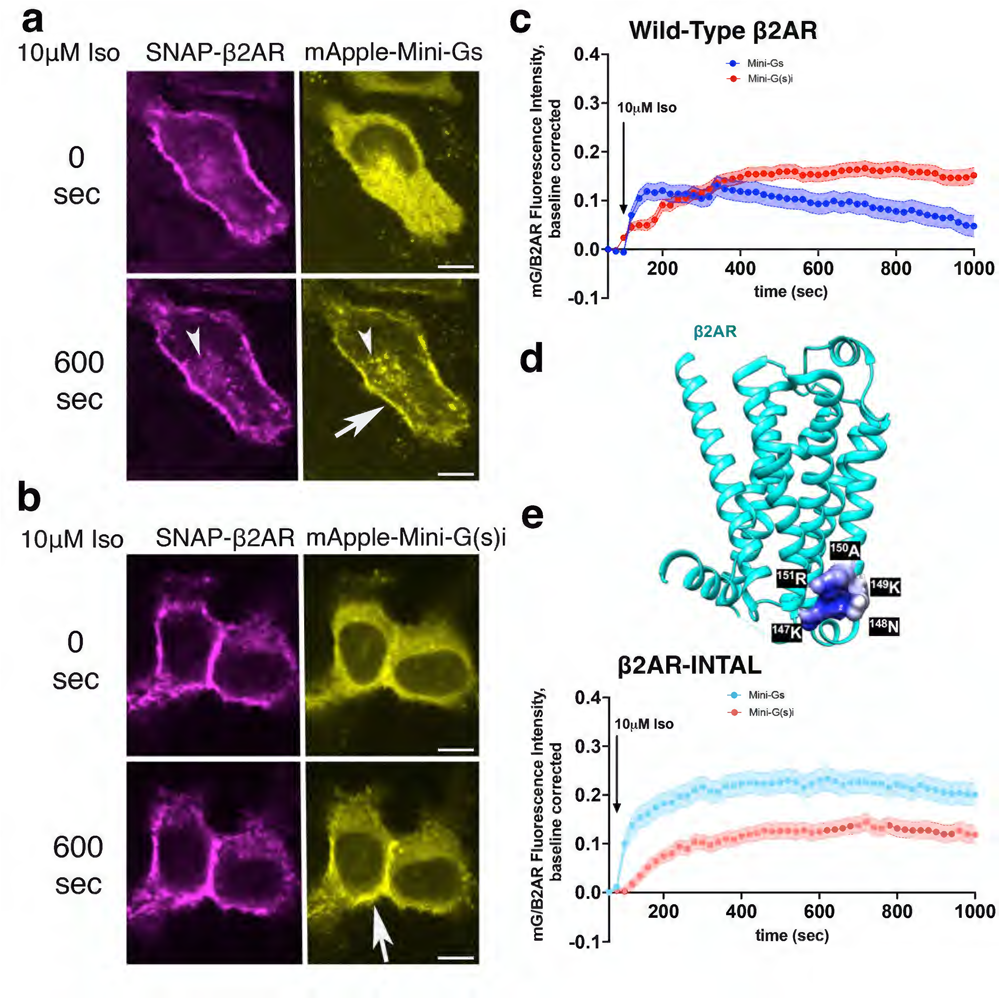
β2AR mutant lacking PI(4,5)P2 binding sites recruits Mini-G(s)i less efficiently. (**a**) Confocal images of representative SNAP-tagged β2AR-expressing HeLa cells (left panels) with mApple-Mini-Gs (right panels) expression before and after 10 μM Iso addition. Stimulation with 10 μM Iso results in recruitment of Mini-Gs to active β2AR at the plasma membrane (n= 49 cells, 4 biological replicates). Arrow indicates mApple-Mini-Gs localization at plasma membrane; Scale bar = 10 μm. (**b**) Confocal images of representative SNAP-tagged β2AR-expressing HeLa cells (left panels) with mApple-Mini-G(s)i (right panels) expression before and after 10 μM Iso addition. Stimulation with 10 μM Iso results in the recruitment of Mini-G(s)i to active β2AR at the plasma membrane (n= 34 cells, 4 biological replicates). Arrow indicates recruited mApple-Mini-G(s)i at plasma membrane; Scale bar = 10 μm. (**c**) Quantification of Mini-G recruitments to activated β2AR at the plasma membrane in HeLa cells after 10 μM Iso addition; normalized fluorescence intensity of Mini-G probes at plasma membrane relative to SNAP-tagged-β2AR. Quantifications were baseline corrected. (**d**) Top: β2AR crystal structure (PDB: 4LDO) with predicted PI(4,5)P2 binding motif at residues 147-151 (KNKAR) showing positive electrostatic charge. Bottom: quantification of Mini-Gα recruitments to activated β2AR INTAL at the plasma membrane in HeLa cells after 10 μM Iso addition; normalized fluorescence intensity of Mini-G probes at plasma membrane relative to SNAP-tagged-β2AR. Quantifications were baseline corrected.

### PI(4,5)P2 allosterically modulates GPCR-mediated cAMP signaling

We next asked whether receptor coupling to opposing Gαs and Gαi proteins has any significant effect on modulating the cAMP response. We compared isoproterenol-mediated cAMP response in wild-type β1AR with PI(4,5)P2 binding mutants, β1AR IATAL. We first checked surface expression of wild-type β1AR and β1AR IATAL and found that β1AR IATAL is substantially expressesed lower at the plasma membrane than wild-type β1AR (Figure 5a). Stimulating HeLa cells expressing wild type β1AR and β1AR IATAL with comparable plasma membrane expression, with 10 μM isoproterenol resulted in significantly higher cAMP accumulation in β1AR IATAL (Figure 5b). We then took advantage of our pseudojanin rapamycin system to assess the effect of rapidly and inducibly depleting PI(4,5)P2 on β1AR-mediated cAMP response. Rapamycin itself had no significant effect on isoproterenol-mediated cAMP response in β1AR expressing HeLa cells (Supplementary Figure 9a). However, depleting PI(4,5)P2 in PM-FKBP and mVenus-FRB-pseudojanin expressing HeLa cells resulted in significantly higher cAMP accumulation in response to 10 μM isoproterenol (Figure 5 c and d). Importantly, depleting PI(4,5)P2 had no significant effect on β1AR-mediated cAMP signaling when HeLa cells were stimulated with 10 μM dobutamine (Supplementary Figure 5b). This is because β1AR can only couple to Gαs when stimulated with dobutamine. These results suggest that PI(4,5)P2 can allosterically modulate βARs coupling to opposing Gαs and Gαi on the plasma membrane thus modulating cAMP response in that location.

**Figure 5.**
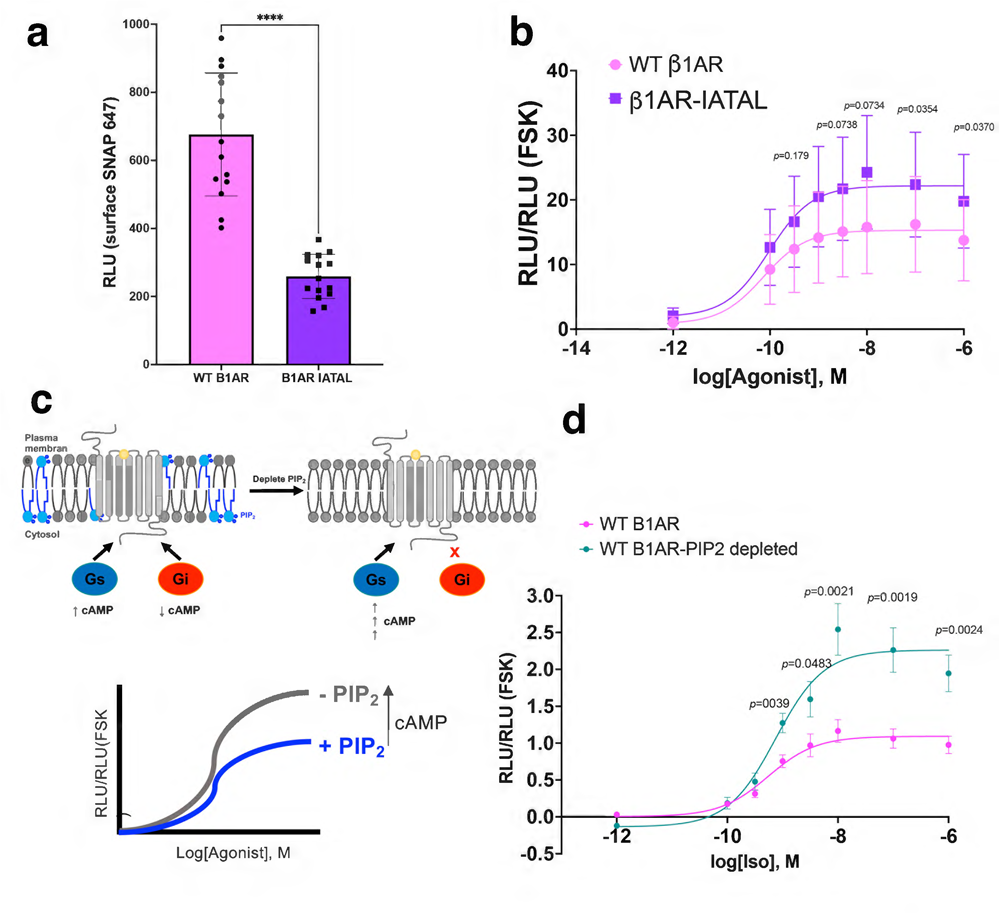
PI(4,5)P2 modulates GPCR-mediated cAMP signaling. (**a**) Surface expression level of HeLa cells expressing wild type β1AR and mutant β1AR IATAL detected by membrane-impermeant SNAP-647 dye (mean +/- SEM, n = 4 biological replicates) (**b**) Forskolin-normalized wild type β1AR and mutant β1AR IATAL-mediated cAMP response with Iso treatment at indicated concentrations in HeLa cells (mean +/- SEM, n= 15 cells, 4 biological replicates). (**c**) Schematic of signaling outputs upon PI(4,5)P2 depletion. Blue colored phospholipids represent PI(4,5)P2 while grey phospholipids represent neutral phospholipids. (**d**) Forskolin-normalized β1AR-mediated cAMP response before and after PI(4,5)P2 depletion upon rapamycin pretreatment (1 μM, 10 min) and at various iso concentrations in HeLa-expressing FKBP-FRB system (mean +/- SEM, n = 4 biological replicates).

## DISCUSSION

The mechanisms determining GPCR-Gα protein selectivity has been studied for decades. Given that there are more GPCRs than Gα proteins, it has been proposed that GPCRs likely evolved to overcome such high scrutiny to couple to specific types of Gα proteins. Various biochemical and structural studies have identified regions within the N-terminal, core, and the C-terminal regions of Gα proteins to play key roles in determining coupling selectivity ^2–5^. One such mechanism proposed that GPCRs binding to the C-terminal domain of Gαs seems to require a more open receptor core, while the Gαi C-terminal domain engages a more closed conformation^36^ and that Gαi-coupled receptors generally reject Gαs proteins due to the bulky C terminus of Gαs^37^. However, a more recent study rejects this model, suggesting that there are no structural barriers to forming a complex between most GPCR–Gα protein pairs and argues the importance of conformational plasticity and highly variable selectivity mechanisms across GPCRs^15^ There are other evolved features that could determine selectivity. These include post-translational modifications, phosphorylation, and potential allostery provided by membrane phospholipids or phosphoinositides. These additional features help GPCRs fine-tune and in some cases, even alter their G protein selectivity patterns^4^.

In this study, we found that the presence of anionic phospholipids, such as PI(4,5)P2, promotes promiscuous coupling to cognate and non-cognate Gα proteins. This non-selective coupling only occurs at the plasma membrane, a membrane compartment enriched in PI(4,5)P2. Furthermore, receptors were able to selectively couple to their cognate Gα proteins at the Golgi membranes, which are enriched in a less negatively charged phospholipids, PI(4)P. These observations support the notion that selectivity can be determined by the lipid environment where receptors reside. A recent NMR study highlights the role of anionic lipids and their interactions with basic residues on the receptor to stabilize and favor a fully active receptor conformation^12^. Our data also supports this model and suggests that PI(4,5)P2 promotes the promiscuous coupling by priming and perhaps stabilizing a fully active receptor conformation that allows receptor coupling to other non-cognate G protein subtypes. This model is further supported by our data showing that dobutamine, a β1AR partial agonist, can only promote Gαs coupling regardless of the membrane lipid environment. This is presumably due to the inability of a partial agonist to stabilize a fully active conformation of β1AR. This priming effect is similar to the effect proposed for G proteins synergistically priming receptor coupling to other non-cognate G proteins^13^.

We also tested whether this promiscuous coupling affects receptor activity. We found that depleting PI(4,5)P2 at the plasma membrane or mutating the predicted PI(4,5)P2 binding sites on exemplary class A GPCR, β1AR, results in enhanced β1AR-mediated cAMP response. Importantly, dual coupling and changes in cAMP levels were only observed when receptors were activated by full agonists (Iso and Epi) and not a partial agonist such as dobutamine. These data further suggest that a fully active conformation of GPCR is required to promote promiscuous coupling. Whether βARs coupling to its non-cognate Gαi protein promotes nucleotide exchange and activation of Gαi protein is unclear. We believed our data support a model where receptor coupling to non-cognate Gα proteins can nevertheless fine-tune receptor-mediated cAMP signaling at the plasma membrane by occupying the βAR-G protein interface and presumably inhibiting nucleotide exchange on the cognate Gαs protein.

Here, we tested two exemplary GPCRs, β1AR and β2AR, that where either shown to contain the PI(4,5)P2 interacting residues^11^, or had high sequence conservation of similar residues at TM4 (Supplementary Figure 7). Whether a similar sequence conservation exists for other Gαs, Gαi and/or Gαq-coupled receptor and whether it is required for promiscuous coupling needs to be determined. Moreover, compartmentalized GPCR signaling has been reported for several other receptors at various subcellular organelles, such as the endosome and the Golgi membranes. It would be important to know whether such coupling selectivity exist for other reported Gαi or Gαq-coupled GPCRs at these subcellular locations^17, 29, 38, 39^.

## Materials and methods

### cDNA constructs/plasmids

#### βAR constructs

Signal Sequence-Snap-tagged β1AR was created by amplifying β1AR from Flag-β1AR 5′-GCTG GGT CTT GGA TCC AAC GAT GAT GAC GCC GGC -3′; 5′- AT AGA ATA GGG CCC TCT AGA GCC CTA CAC CTT GGA TTC CG -3′ primers and inserted into the Snap vector using BamHI and XbaI. Mutant β1AR IATAL and mutant β2AR INTAL were created by site directed mutagenesis using 5’- CG ATT GCT ACA GCT CTG GGC CTC GTG TGC ACC GT -3’; 5’- GCC CAG AGC TGT AGC AAT CGT CAG CAG GCT CTG GTA GCG -3’ primers and 5’-CC ATT AAT ACA GCA TTA GTG ATC ATT CTG ATG GTG TGG ATT G -3’; 5’- AC TAA TGC TGT ATT AAT GGT CAG CAG GCT CTG GTA C -3’ primers, respectively. Flag-β1AR-NanoLuc was a gift from the Lambert lab.

#### Mini-G probes

pmVenus-Mini-Gs, pmVenus-Mini-G(s)I constructs were gifts from Dr. Nevin Lambert. pmVenus-Mini-G(s)i EDGE and pmApple-Mini-G(s)i EDGE were created from pmVenus-Mini-G(s)i and pmApple-Mini-G(s)i by site directed mutagenesis using 5’- A AAA CAG CTC CAG AAA GAC GGG CAA GTG TAC AGG G -3’; 5’- T CTA TGA GTG GCC CTG TAC ACT TGC CCG TCT TTC TGG -3’ primers for the first round of mutation (KDKQ → KDGQ), and 5’- GAA AAA CAG CTC CAG GAA GAC GGG GAA GTG -3’; 5’- AGT GGC CCT GTA CAC TTC CCC GTC TTC CTG -3’ primers for the second round of mutation (KDGQ → EDGE). C-terminal domain swapped Mini-G probes were created using site directed mutagenesis using 5’- G AAT CTT CGG CAA TAC GGG CTT TTC TAA AAG CTT CGA A -3’; 5’- CG AAG CTT TTA GAA AAG CCC GTA TTG CCG AAG ATT CA -3 for the first round of mutation (DCGLF → QYGLF) and 5’- CTT CGG CAA TAC GAG CTT CTC TAA AAG CTT CGA AT -3’; 5’- TT TTA GAG AAG CTC GTA TTG CCG AAG ATT CAT TTT GA -3 for the second round of mutation (QYGLF → QYELL) of the Gs C-terminal domain swap, and using 5’- G CAT CTC AGG GAC TGT GAG CTG CTC TAA AAG CTT CG -3’; 5’- C GAA GCT TTT AGA GCA GCT CAC AGT CCC TGA GAT GCA -3 for the first round of mutation (QYELL → DCELL) and 5’- TC AGG GAC TGT GGG CTG TTC TAA AAG CTT C -3’; 5’- AGC TTT TAG AAC AGC CCA CAG TCC CTG AGA TGC -3 for the second round of mutation (DCELL → DCGLF) of the Gi C-terminal domain swap.

#### Rapamycin heterodimerization constructs

Lyn-2xFKBP-Myc was used in a previous study ^23^. pmVenus-FRB-Pseudojanin (PJ) construct was created from mRFP1-1xFKBP-PJ (Hammond et al., 2012; Addgene) by amplifying PJ using 5’- GT GCT GGT GGA GGA TCC AGT GCT GGT GGT AGT GCT - 3’; 5’- TTA TCT AGA TCC GGT GGA TCC TCA AGA AAC GGA GGC GAT G -3’ primers and inserted into the pmVenus-FRB-C1 vector using BamHI.

### Cell culture and transfections

HeLa cells (purchased from ATCC as authenticated lines CCL-2) were grown in Dulbecco’s minimal essential medium supplemented with 10% fetal bovine serum (FBS) without antibiotics. Cell cultures were free of mycoplasma contamination. Transfections were performed using Transit-LT1 transfection reagent (Mirus Bio) at a 1:3 DNA to transfection reagent ratio. Snap-tagged human β1AR and β2AR constructs were labeled with Snap-cell 647 SiR (New England Biolabs, S9102S) at 1:1000 concentration for 20 min at 37°C in DMEM without phenol red supplemented with 30 mM HEPES, pH 7.4, washed with the same media (thrice for one minute, room temperature), and imaged in the same media.

### Live-cell confocal imaging

Live-cell imaging was carried out using a Nikon spinning disk confocal microscope with a ×60, 1.4 numerical aperture, oil objective and a CO2 and 37°C temperature-controlled incubator. A 488, 568 nm and 640 Voltran was used as light sources for imaging mVenus/GFP, mApple, and Snap-647 signals, respectively. Cells expressing both Snap-tagged receptor (1 μg for both wildtype β1AR and β2AR, 4 μg for mutant β1AR IATAL and 2 μg mutant β2AR INTAL) and the indicated Mini-G probe (100 ng) were plated onto glass coverslips. Receptors were surface labeled by addition of Snap-Cell 647 SiR (1:1000, New England Biolabs) to the media for 20 min, as described previously. Indicated agonists (isoproterenol, epinephrine, or dobutamine all from Sigma) were added and cells were imaged every 20 s for 20 min in DMEM without phenol red supplemented with 30 mM HEPES, pH 7.4. rapamycin (Sigma) was incubated for 10 min before cells were stimulated with 10 μM isoproterenol. Time-lapse images were acquired with a CMOS camera (Photometrics) driven by Nikon Imaging Software (NS Elements).

### Image analysis and statistical analysis

Images were saved as 16-bit TIFF files. Quantitative image analysis was carried out on unprocessed images using ImageJ software (http://rsb.info.nih.gov/ij). For measuring kinetics of Mini-G probe recruitment at the plasma and Golgi membranes over time in confocal images, analyses were performed on unprocessed TIFF images using a previously published scripts written in MATLAB, available through open access on zenoob ^40–42^. Briefly, the plasma membrane and Golgi regions were selected and a mask of labeled receptor (using Snap label) or Golgi marker were generated by thresholding the receptor or the Golgi marker signal within the selected region. The average fluorescence intensity of Mini-G was measured within the masked region and outside of the masked region, before and after addition of agonists. Values were normalized by calculating the percent relative to the maximum value, then baseline corrected using GraphPad Prism v.9.5.1 (GraphPad) software so that the first value of each condition was set to 0.

### Luminescence-based cAMP assay

HeLa cells were transfected with either wildtype β1AR, wildtype, β2AR, mutant β1AR IATAL, or mutant β2AR INTAL and a plasmid encoding a cyclic-permuted luciferase reporter construct (pGloSensor-20F, Promega) and luminescence values were measured, as described previously ^43^. Briefly, cells were plated in 96-well dishes (∼100,000 cells per well) in DMEM without phenol red/no serum and equilibrated to 37°C in the SpectraMax plate reader and luminescence was measured every 1.5 min for 20 min. Software was used to calculate integrated luminescence intensity and background subtraction. In rapamycin heterodimerization experiments, cells were pre-incubated with 1 μM rapamycin (Sigma) for 10 min. 10 μM forskolin (Sigma) was used as a reference value in each multi-well plate and for each experimental condition. The average luminescence value (measured across duplicate wells) was normalized to the maximum luminescence value measured in the presence of 10 μM forskolin. For rapamycin-treated cells, the average luminescence value was normalized to the maximum luminescence value measured in the presence of 10 μM forskolin and 1 μM rapamycin. Concentration–response curves were fitted to a three-parameter binding logistic model using GraphPad Prism v.9.5.1 (GraphPad).

### Western blotting

HeLa cells were lysed in extraction buffer (0.2% Triton X-100, 50 mM NaCl, 5 mM EDTA, 50 mM Tris at pH 7.4 and cOmplete EDTA-free Protease Inhibitor Cocktail [Roche]). After agitation at 4°C for 30 min, supernatants of samples were collected after centrifuging at 15,000 × rpm for 10 min at 4°C. Supernatants were mixed with SDS sample buffer for the protein denaturation. The proteins were resolved by SDS-PAGE and transferred to PVDF membrane and blotted for anti-SNAP (1:1000) or GAPDH (1:10,000) antibodies to detect wildtype and mutant Snap-labeled β1AR and β2AR and GAPDH expression by horseradish-peroxidase-conjugated rabbit IgG, sheep anti-mouse and rabbit IgG (1:10,000 Amersham Biosciences), and SuperSignal extended duration detection reagent (Pierce).

### BRET, luminescence measurements

HeLa cells were transiently transfected with Flag-β1AR-NanoLuc, pmVenus-Mini-Gs, pmVenus-Mini-G(s)i, pmApple-Mini-Gs, pmApple-Mini-G(s)i at a 1:4 donor: acceptor ratio (receptor: Mini-G). After a 24-h incubation, cells were washed thrice with 1× HBSS, harvested by trypsinization, and transferred to opaque black 96-well plates. Kinetic BRET and luminescence time course measurements were made using a SpectraMax plate reader. Furimazine (NanoGlo; 1:1000, Promega) was added to all wells immediately prior to making measurements with Nanoluc. Raw BRET signals were calculated as the emission intensity at 530 nm divided by the emission intensity at 480 nm. Net BRET signals were calculated as the raw BRET signal measured from cells expressing Flag-β1AR-NanoLuc and mVenus-Mini-G minus the raw BRET signal measured from cells expressing Flag-β1AR-NanoLuc and mApple-Mini-G. Heatmaps represent ΔBRETmax (Figure 1e).

### Sequence alignments and receptor structures

All sequence alignments in Supplementary information (Supp figs. 2 and 4) were performed with Clustal Omega Multiple Sequence Alignment (https://www.ebi.ac.uk/Tools/msa/clustalo/). Sequences were extracted from UniProt (https://www.uniprot.org/). Structures in Figures 2 and 4 correspond to a previous publication (Xu et al., 2021), and can be found in Protein Data Bank (PDB: 7BTS and 4LDO, β1AR and β2AR, respectively). Figures were generated using UCSF Chimera v.1.17.3.

## Data availability

All data generated and analyzed during this study are included in this manuscript as figures, and supplementary figures.

## Acknowledgments

We thank members of the Irannejad lab, especially Dr. Quynh N. Mai for assistance, advice, and valuable discussion. We would like to thank Members of the Center for Advanced Microscopy, Drs. Larsen, Kari Herrington and So Yeon Kim for assistance with the microscopy experiments. These studies were supported by the Ford Foundation Fellowship to E.H.H., the National Institute on General Medicine (GM133521) to R.I. and (GM138992) to A.M.

---

**Supplementary figure 1.**
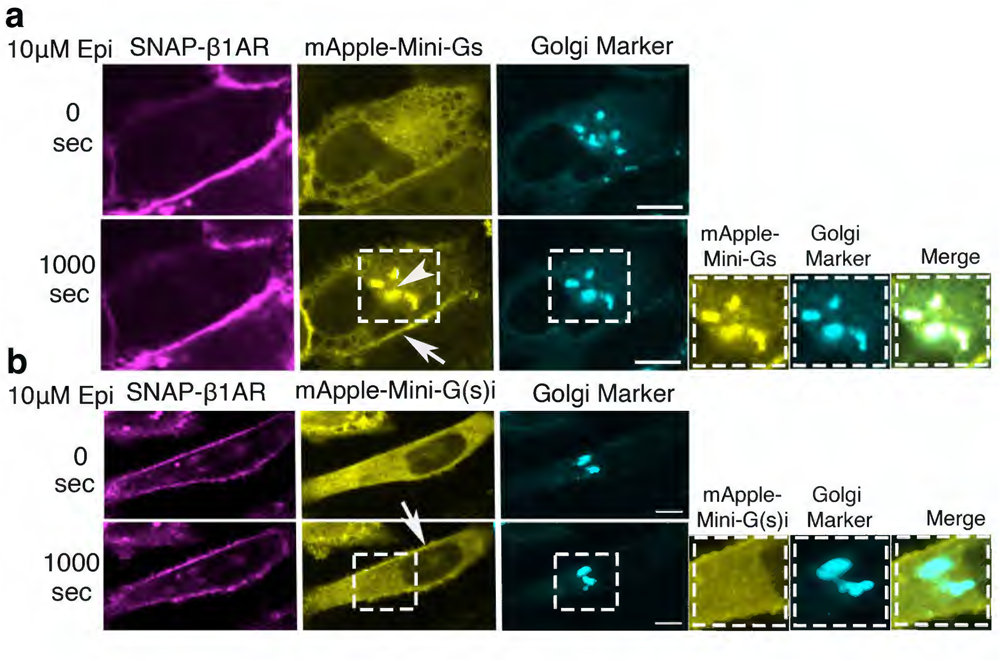
β1AR couples to Gαs proteins only at the Golgi membrane upon epinephrine (Epi) stimulation, as detected by Mini-Gs recruitment. **(a)** Confocal images of representative SNAP-tagged β1AR-expressing HeLa cells (left panels) with mApple-Mini-Gs (middle panels) before and after 10 μM Epi addition. Stimulation with 10 μM Epi results in recruitment of Mini-Gs to active β1AR at the Golgi apparatus, indicated by the Golgi marker (right panels) in HeLa cells (n= 26 cells, 3 biological replicates). Arrow and arrowhead indicate Mini-Gs recruitment at the plasma membrane and Golgi, respectively; Scale bar = 10 μm. Right panels show zoomed images of insets for Mini-Gs, Golgi marker, and merge. **(b)** Confocal images of representative SNAP-tagged β1AR-expressing HeLa cells (left panels) with mApple-Mini-G(s)i (middle panels) before and after 10 μM Epi addition. Stimulation with 10 μM Epi does not result in the recruitment of Mini-G(s)i to active β1AR at the Golgi membranes (n= 14 cells, 3 biological replicates). Arrow indicates recruited Mini-G(s)i at the plasma membrane only; Scale bar = 10 μm. Right panels show zoomed images of insets for Mini-G(s)i, Golgi marker, and merge.

**Supplementary figure 2.**
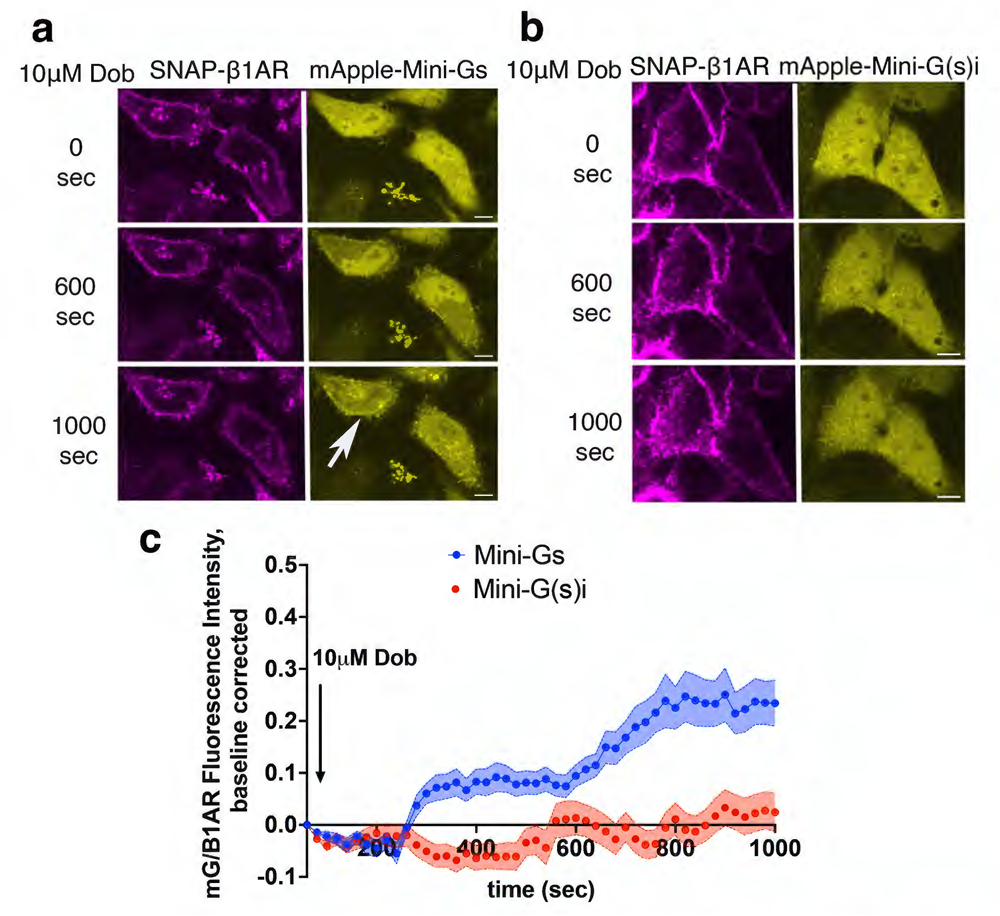
β1AR couples to Gαs proteins only at the plasma membrane and Golgi membrane upon dobutamine (Dob) stimulation, as detected by Mini-Gs recruitment. **(a)** Confocal images of representative SNAP-tagged β1AR-expressing HeLa cells with mApple-Mini-Gs expression before and after 10 μM Dob addition. Stimulation with 10 μM Dob results in recruitment of Mini-Gs (n= 17 cells, one biological replicate) to active β1AR at the plasma membrane and Golgi in HeLa cells. Arrow and arrowhead indicate Mini-Gs recruitment to the plasma membrane and the Golgi, respectively; Scale bar = 10 μm. (**b**) Confocal images of representative Snap-tagged β1AR-expressing HeLa cells and mApple-Mini-G(s)i expression before and after 10 μM Dob addition. Stimulation with 10 μM Dob does not result in recruitment of Mini-G(s)i (n= 15 cells, one biological replicate) to active β1AR at the plasma membrane or Golgi in HeLa cells. Scale bar = 10 μm. **(c)** Quantification of Mini-Gα recruitments to activated β1AR at the plasma membrane in HeLa cells after 10 μM Dob addition; normalized fluorescence intensity of Mini-G probes at plasma membrane relative to SNAP-tagged-β1AR. Quantifications were baseline corrected after addition of Iso.

**Supplementary figure 3.**
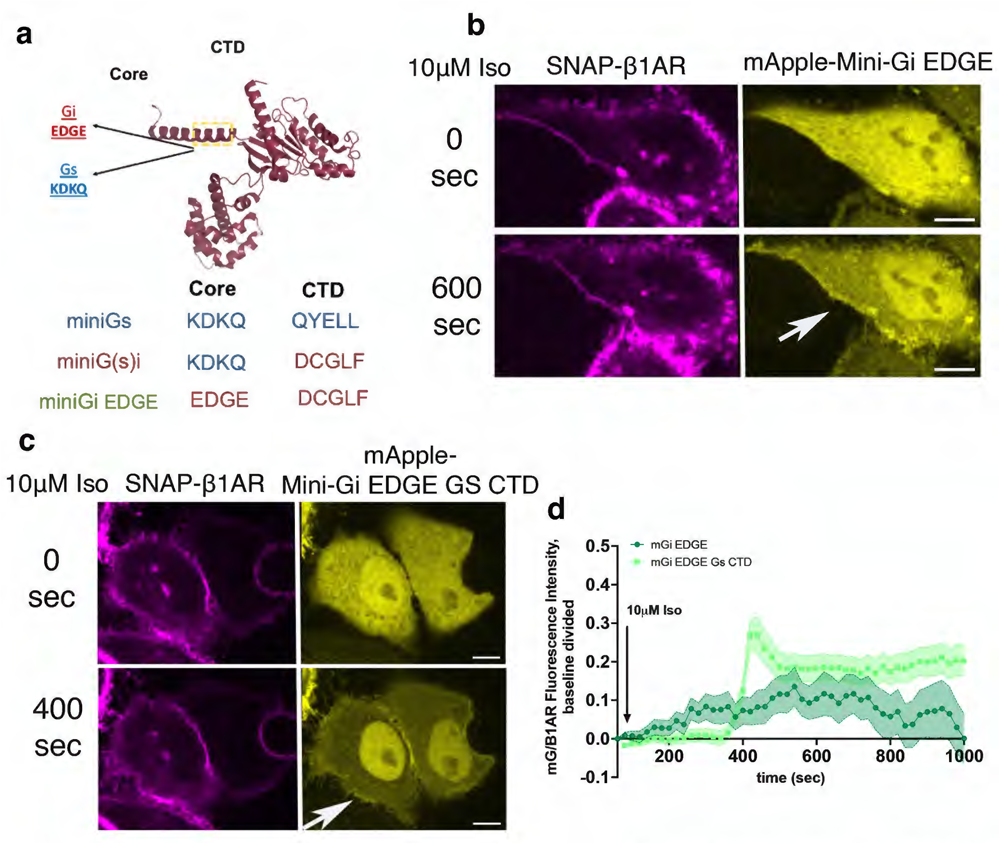
N-terminal and core region of Gαs promote efficient coupling to activated β1AR upon isoproterenol stimulation. **(a)** The N-terminal core region of Mini-G(s)i was mutated from its positive sequence (KDKQ) to negative sequence (EDGE) to create Gαi-like biosensor, Mini-G(s)i EDGE. **(b)** Confocal images of representative SNAP-tagged β1AR-expressing HeLa cells (left panels) with mApple-Mini-G(s)i EDGE (right panels) expression before and after 10 μM Iso addition. Stimulation with 10 μM Iso results in recruitment of Mini-G(s)i EDGE (n= 14 cells, 3 biological replicates) to active β1AR at the plasma membrane in HeLa cells. Arrows indicate recruited Mini-G(s)i EDGE at plasma membrane; Scale bar = 10 μm. **(c)** Confocal images of representative SNAP-tagged β1AR-expressing HeLa cells (left panels) with mApple-Mini-G(s)i EDGE with Gs CTD (right panels) before and after 10 μM Iso addition. Stimulation with 10 μM Iso results in recruitment of Mini-Gi EDGE with Gαs CTD (n= 24 cells, 1 biological replicate) to active β1AR at the plasma membrane in HeLa cells. Arrows indicate recruited Mini-G(s)i EDGE with Gαs CTD at plasma membrane; Scale bar = 10 μm. **(d)** Quantification of Mini-Gα recruitments to activated β1AR at the plasma membrane in HeLa cells after 10 μM Iso addition; normalized fluorescence intensity of Mini-G(s)i EDGE and Mini-G(s)i EDGE with Gαs CTD probes at plasma membrane relative to SNAP-tagged-β1AR. Quantifications were baseline corrected.

**Supplementary figure 4.**
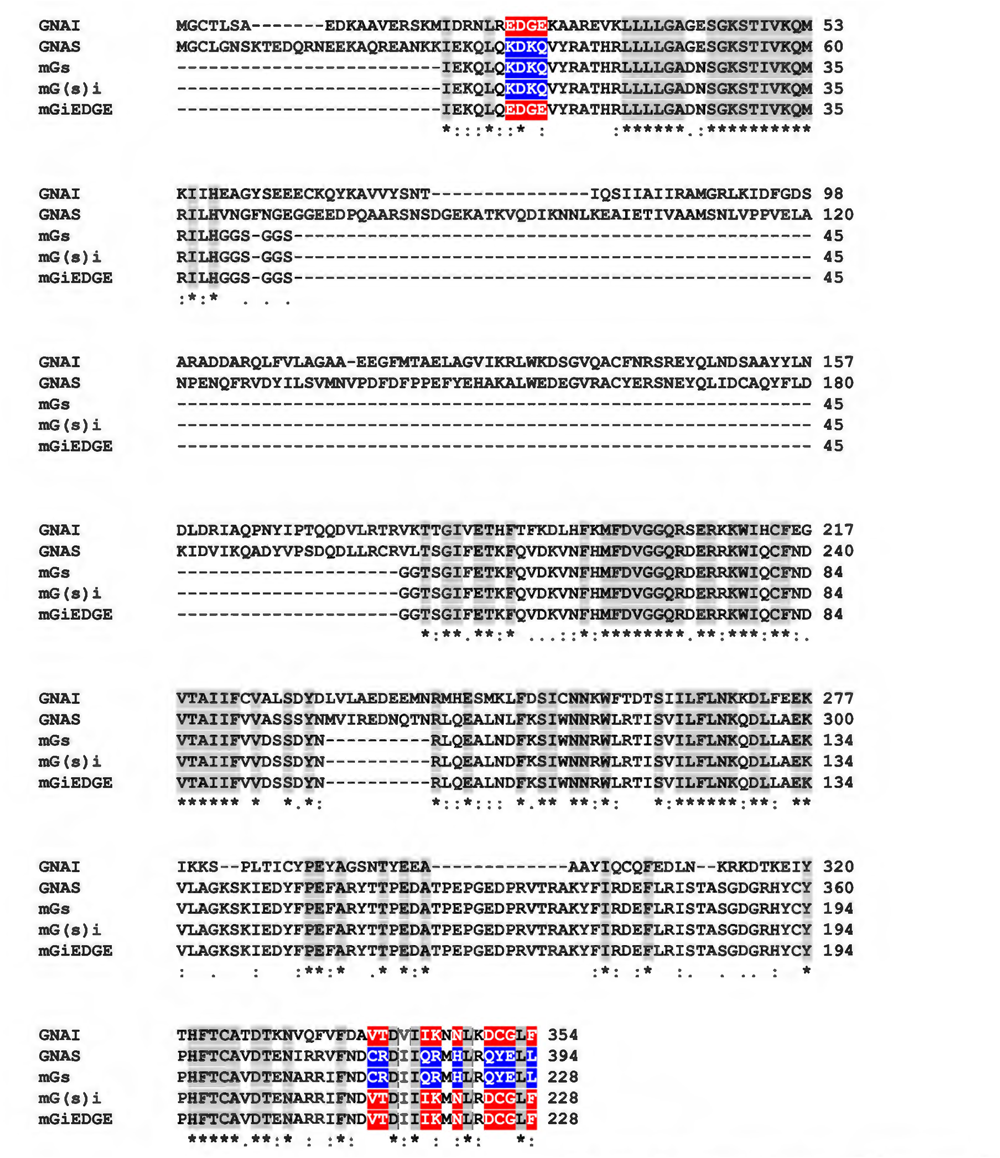
Protein sequence alignment for human GNAI, human GNAS, Mini-Gs, Mini-G(s)i, and Mini-G(s)i EDGE. The alignment shows the segments from GNAS and GNAI that were excluded in the Mini-G probes as well as the segments which were swapped to make the Mini-G(s)i chimera. Segments in red represent the N-terminal core and the C-terminal regions that originate from GNAI. Segments in blue represent the N-terminal core and C-terminal regions that originate from GNAS. Segments with colons (:) represent sequences with similar amino acid properties such as electrostatic charge, hydrophobicity, and/or side chain size. Alignments were performed using Clustal Omega Multiple Sequence Alignment.

**Supplementary figure 5.**
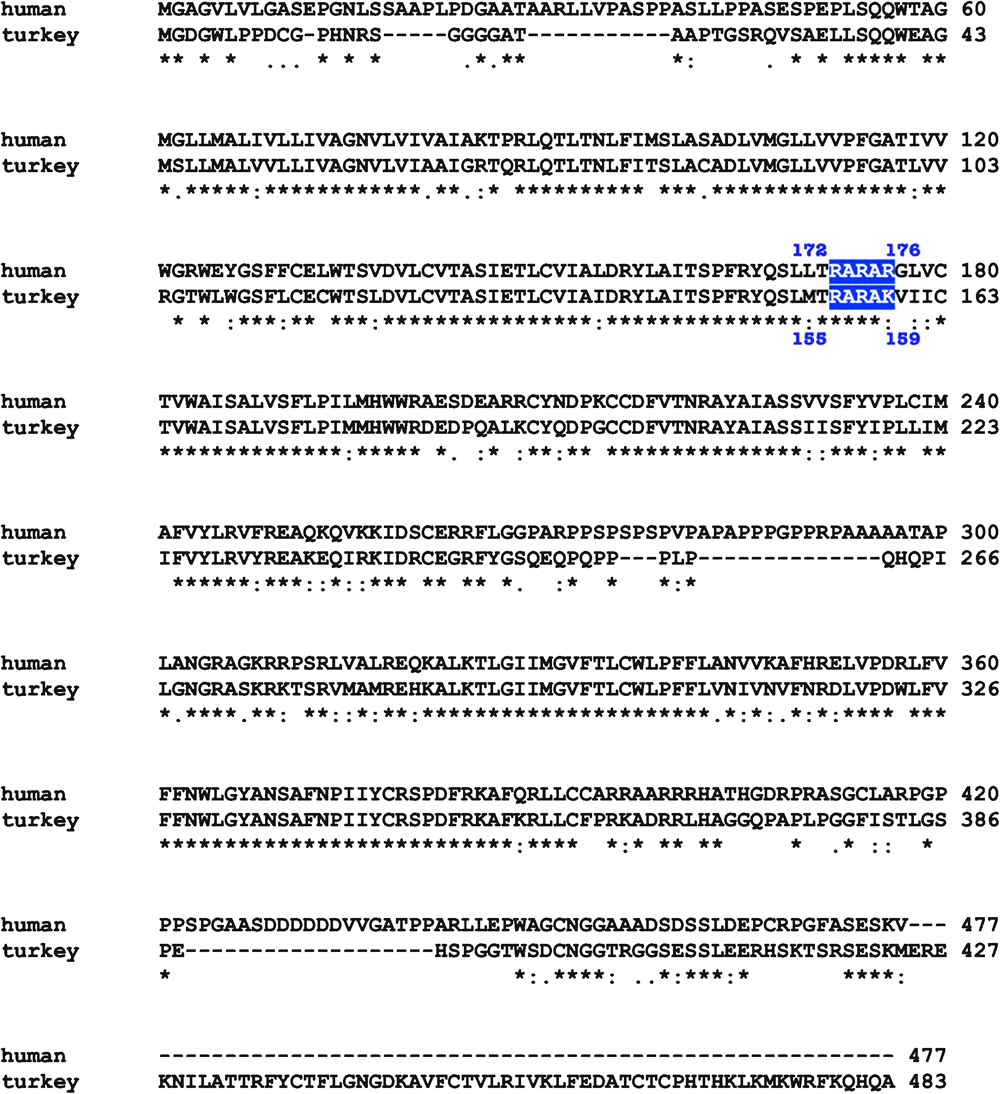
Protein sequence alignments of human and turkey β1AR. Segments in blue represent PI(4,5)P2 binding region previously identified for turkey β1AR. Alignments were performed using Clustal Omega Multiple Sequence Alignment.

**Supplementary figure 6.**
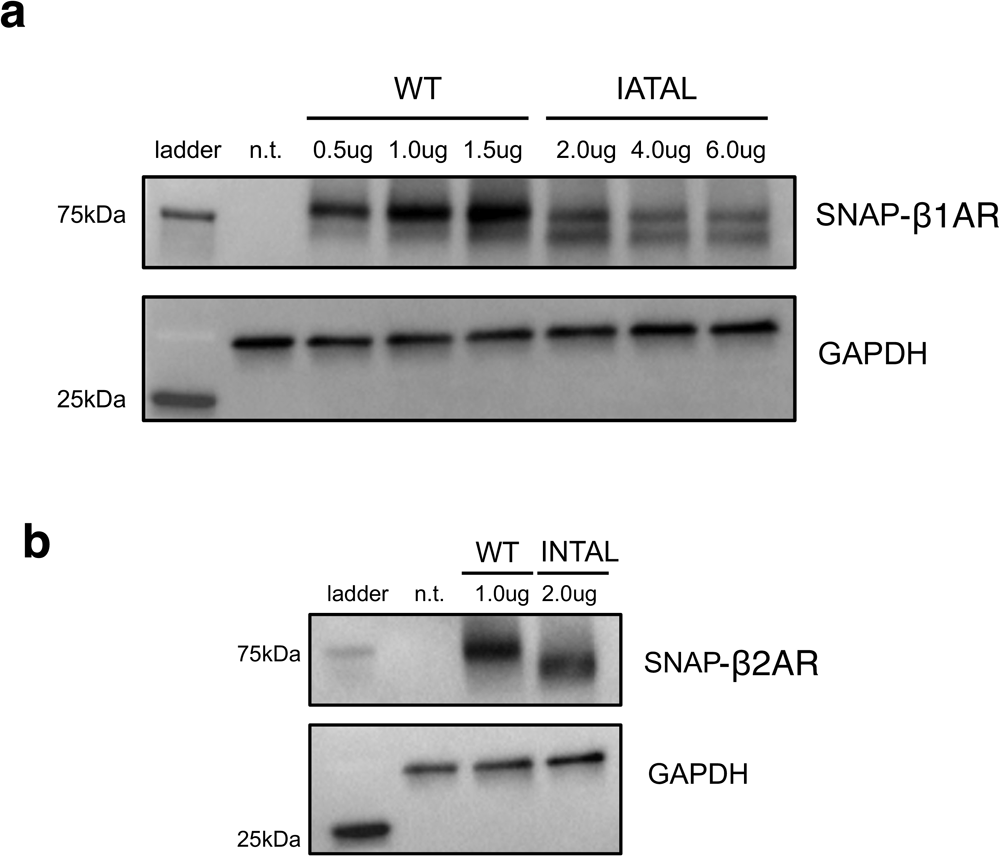
Protein expression levels of wild type βAR compared to PI(4,5)P2 binding mutants. (**a**) Representative western blots of HeLa cells expressing wild type SNAP-tagged-β1AR and mutant SNAP-tagged-β1AR IATAL, detected by anti-SNAP antibody. Lanes containing 0.5 μg WT β1AR and 2.0 μg mutant β1AR IATAL DNA transfection in HeLa cells showed the closest expression levels. (**b**) Representative western blots of HeLa cells expressing wild type SNAP-tagged-β2AR and mutant SNAP-tagged-β2AR INTAL, detected by anti-SNAP antibody. Lanes containing 1.0 μg WT SNAP-tagged-β2AR and 2.0 μg mutant SNAP-tagged-β2AR INTAL DNA transfection in HeLa cells showed the closest expression levels. GAPDH antibody was used as a loading control.

**Supplementary figure 7.**
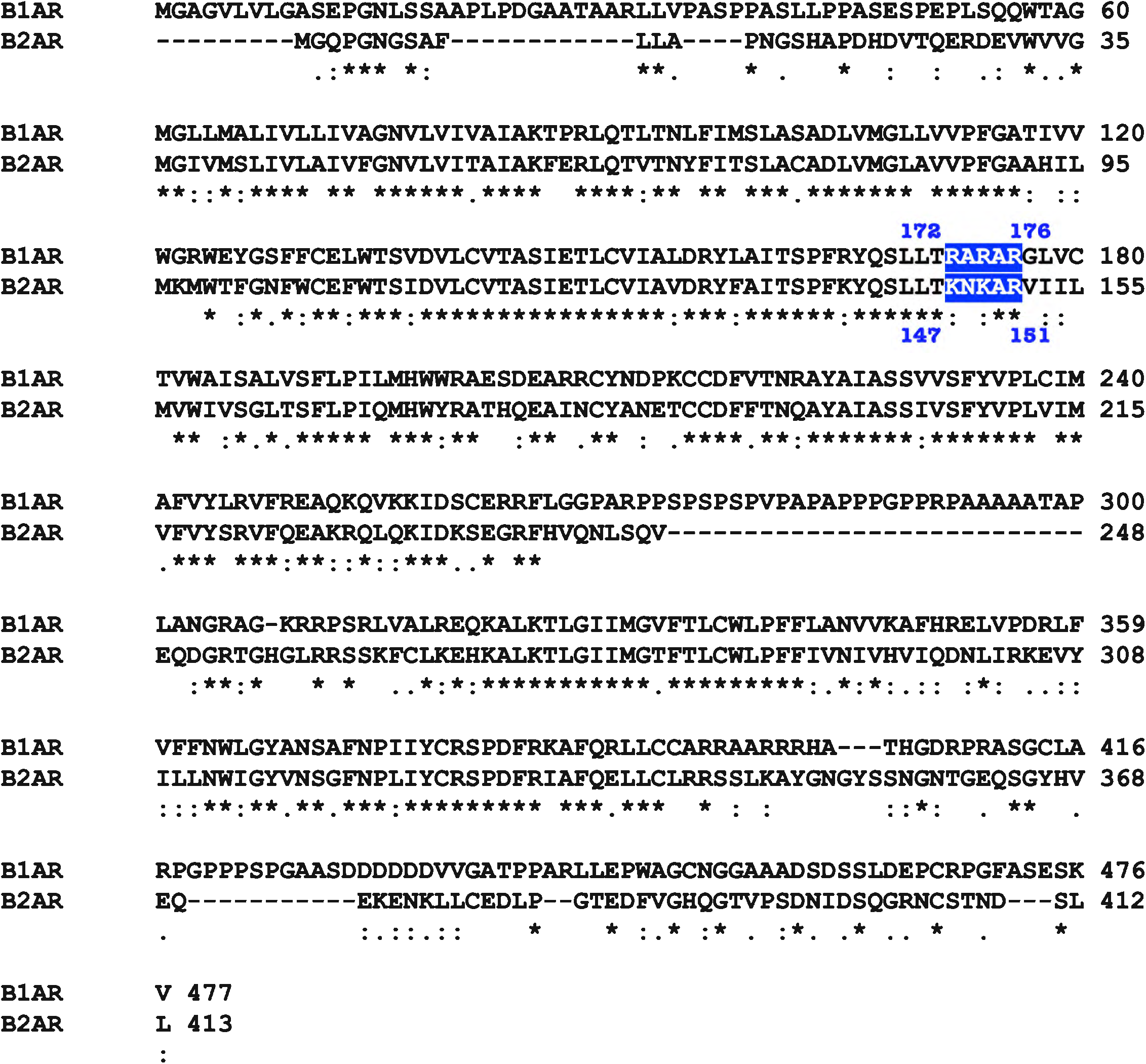
Protein sequence alignment for human β1AR and β2AR. Segments in blue represent PI(4,5)P2 binding regions in human β1AR and human β2AR. Alignments were performed using Clustal Omega Multiple Sequence Alignment.

**Supplementary figure 8.**
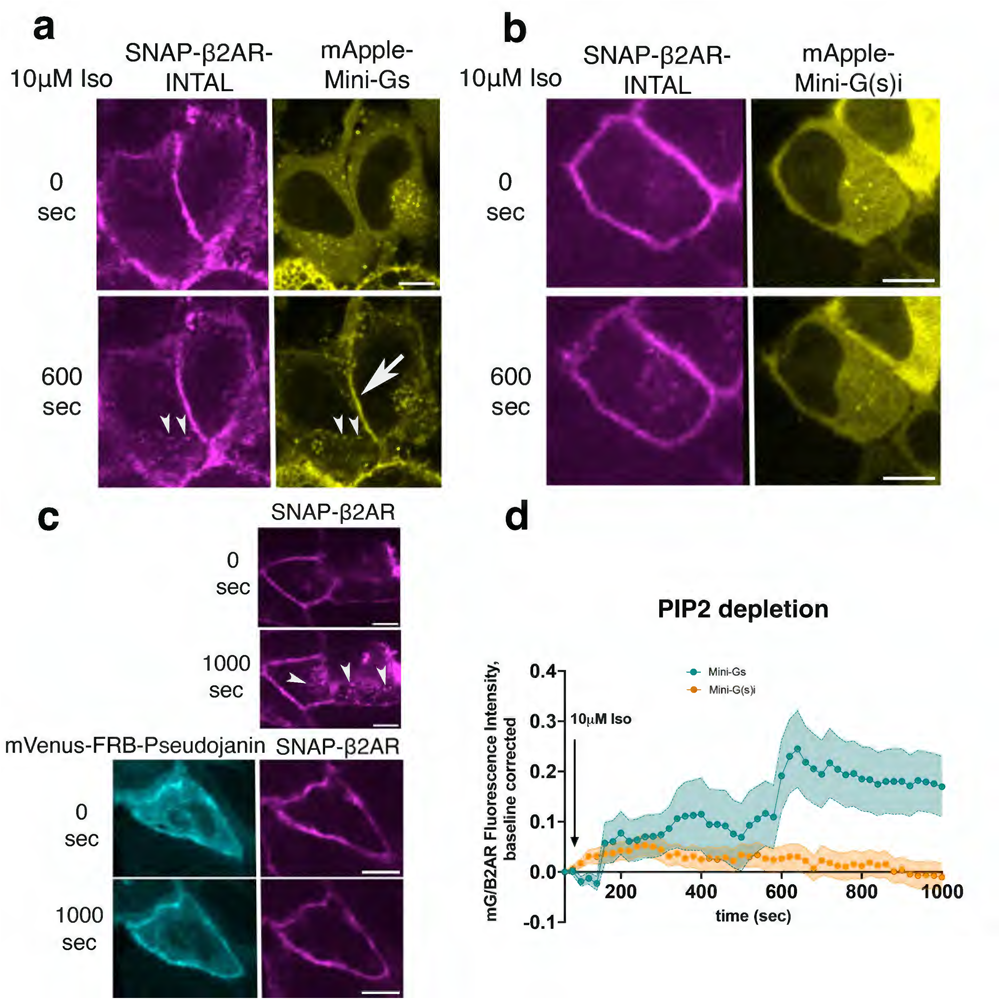
β2AR PI(4,5)P2 binding mutant modestly recruits Mini-G(s)i. (**a**) Confocal images of representative SNAP-tagged β2AR INTAL expressing HeLa cells (left panels) with mApple-Mini-Gs (right panels) before and after 10 μM Iso addition (n= 64 cells, 4 biological replicates). Arrow indicates Mini-Gs recruitment at plasma membrane; arrowheads indicate internalized mutant β2AR INTAL along with Mini-Gs; Scale bar = 10 μm. (**b**) Confocal images of representative SNAP-tagged mutant β2AR INTAL expressing HeLa cells (left panels) with mApple-Mini-G(s)i (right panels) before and after 10 μM Iso addition (n= 54 cells, 4 biological replicates). Scale bar = 10 μm. (**c**) Top: confocal images of WT SNAP-tagged-β2AR. Arrowheads indicate internalized WT β2AR at similar levels as mutant β2AR INTAL. Bottom: confocal images of PI(4,5)P2 depletion using the FKBP-FRB system with WT SNAP-tagged-β2AR (right panels). mVenus-FRB-Pseudojanin is recruited to the plasma membrane upon addition of 1 μM Rapamycin (left panels). Internalization of WT β2AR is not observed when PI(4,5)P2 is depleted. (**d**) Quantification of Mini-G recruitments to activated β2AR at the plasma membrane upon PI(4,5)P2 depletion in HeLa cells after 10 μM Iso addition; normalized fluorescence intensity of Mini-G probes at plasma membrane relative to SNAP-tagged-β2AR. Quantifications were baseline corrected.

**Supplementary figure 9.**
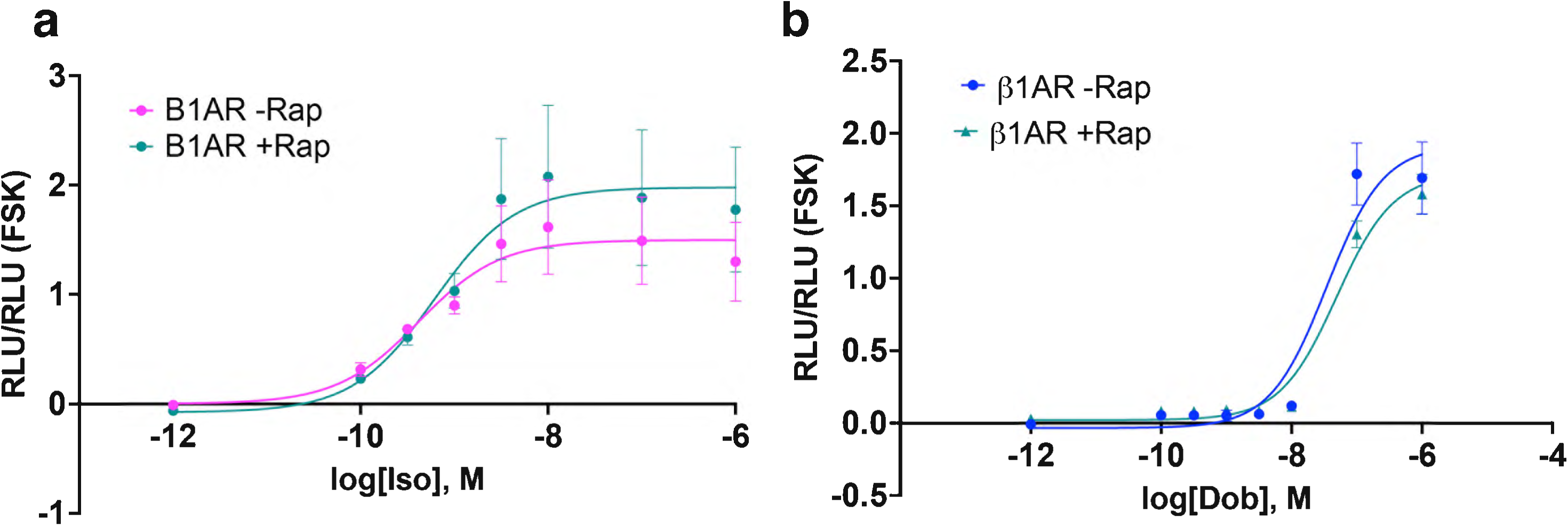
PI(4,5)P2 depletion had no effect on dobutamine-mediated cAMP signaling in β1AR expressing HeLa cells. (**a**) Forskolin-normalized β1AR-mediated cAMP response with and without Rapamycin pretreatment (1 μM, 10 min) in HeLa cells not expressing FKBP-FRB system (mean +/- SEM, n = 2 biological replicates). (**b**) Forskolin-normalized β1AR-mediated cAMP response before and after PI(4,5)P2 depletion upon rapamycin pretreatment (1 μM, 10 min) and at various Dob concentrations in HeLa-expressing FKBP-FRB system (mean +/- SEM, n = 2 biological replicates).

